# Changes in the potential distribution of the guava fruit fly *Anastrepha striata* (Diptera, Tephritidae) during the current and possible future climate scenarios in Colombia

**DOI:** 10.1101/2020.09.26.315143

**Authors:** Eduardo Amat, Mariano Altamiranda-Saavedra, Nelson A Canal, Luz M Gómez-Piñerez

## Abstract

Climate change has affected the geographical distributions of most species worldwide; in particular, insects of economic importance inhabiting tropical regions have been impacted. Current and future predictions of change in geographic distribution are frequently included in species distribution models (SDMs). The potential spatial distributions of the fruit fly *Anastrepha striata* Schiner (the main species of agricultural importance in guava crops) under current and possible future scenarios in Colombia were modeled, and the establishment risk was assessed for each guava-producing municipality in the country. The SDMs were developed using 221 geographical records in conjunctuin with nine scenopoetic variables. The model for current climate conditions indicated an extensive suitable area for the establishment of *A. striata* in the Andean region, smaller areas in the Caribbean and Pacific, and almost no areas in the Orinoquia and Amazonian regions. A brief discussion regarding the area suitability for the fly is offered. The expansion of the suitable area was observed in all future scenarios; moreover, this effect was more pronounced in the Amazonian region. The Colombian guava-producing municipalities were classified according to the degree of vulnerability to the fly establishment as follows: 42 were high-risk, 16 were intermediate-risk, and 17 were low-risk. The implementation of future integrated management plans must include optimal spatial data and must consider environmental aspects, such as those suggested by the models presented here. Control decisions should aim to mitigate the positive relationship between global warming and the increase in the dispersal area of the fruit fly.

## Introduction

Climate change is expected to cause shifts in the geographical distribution of species as a result of the rearrangement of climate zones (Beever et al. 2011). Hence, the magnitude of the associated impacts is projected to be higher in some regions than in others. The Latin American and Caribbean region is one of the most vulnerable areas to climate change; most of the species living there are endemic or restricted to a specific tropical ecosystem (CEPAL 2015). Consequently, they are more susceptible to the effects of global warming because of their particular physiology and phenological qualities, which are typically adapted to narrow ecological niches (Sheldon 2019). It is anticipated that poikilothermic organisms such as insects, whose body temperature varies according to surrounding weather, will be strongly influenced by a volatile climate (Régnière et al. 2012). Temperature, precipitation, and other climatic parameters can directly affect the ecological interactions of insect pests; for instance, the increase in heat in the tropics allows species to colonize higher elevations and extend their geographical distributions upslope (Freeman et al. 2018). Indeed, climate warming resulting from increasing levels of greenhouse gases in the Earth’s atmosphere could have a significant and highly uncertain impact on the development, distribution, and population density of agricultural insect pests (Lehmann et al. 2020).

Predictions of geographical distribution changes relating to global warming are frequently included in species distribution models (SDMs) (Guisan and Zimmermann 2000). These models use associations between environmental variables, such as temperature, precipitation and geographical records of species to identify the environmental conditions under which reproductive populations can be established (Peterson et al. 2011); SDMs have multiple applications in conservation, ecology, evolution, epidemiology, and invasive species management studies (Peterson 2006). In an agricultureural context, SDMs allow the assessment of the potential dispersal of exotic and invasive species to crops (Villacide and Corley 2003; Beckler et al. 2005; Campo et al. 2011), while also permitting the implementation of control and eradication programs and monitoring of these biological agents. SDMs can also can assist in the selection of cultivable areas and declaration of phytosanitary problem-free-zones (Anderson and Martínez-Meyer 2004; Parra et al. 2004; Phillips et al. 2006; Aluja and Mangan 2008). The advantages of these models make their use appropriate in making decisions to mitigate the effects of insect pests.

One of the most common crop-limiting insects is the guava fruit fly *Anastrepha striata* Schiner, 1868 (Diptera, Tephritidae); it endemic species to the Neotropical region, and it is listed as a quarantine species. *A. striata* has been reported to be associated with thirty-seven host plant species in twenty-three genera and seventeen families; most are in Myrtaceae, which is a primary host taxon (Norrbom 2004; Cruz-López et al. 2015). This fruit fly causes substantial agricultural losses, particularly in guava crops, thoroughout the American continent (Castañeda et al. 2010). It is distributed from the southern USA to Brazil (Hernandez-Ortiz and Aluja 1993), with an altitudinal distribution between 15 and 2,398 meters of elevation (Martinez and Serna 2005; Castañeda et al. 2010). In Colombia, this fruit fly is common and has been systematically collected (Rodriguez Clavijo et al. 2018); its prevalence in this area is typically associated with the hostplant (Gallo-Franco et al. 2017). In particular, the species has been reported on guava crops, turning into a plague with significant negative impacts on fruit production (Insuasty et al. 2007; Martinez-Alava 2007; Castañeda et al. 2010). In Colombia, guava is one of the top five species of economic importance and is a significant crop in Colombian agriculture as an essential product of small and intermediate producers (Agronet 2018). The damage caused by *A. striata* can be devastating; total losses of 90% of the crop have been documented recent decades (Núñez et al. 2004). Management plans, including the potential distribution of fruit flies, have been considered in the United States (Sequeira et al. 2001), Europe (Godefroid et al. 2015), and globally for *Anastrepha obliqua* (Fu et al. 2014). In Colombia, integrated pest management against fruit flies has been proposed by governmental institutions (Instituto Colombiano de Agropecuario - ICA); however, none of these initiatives have included potential distribution or spatial distribution modeling (ICA 2016). We aimed to model the potential distribution of *A. striata* and to assess the establishment risk in Colombia under current and possible future climate change scenarios. The resulting maps and concerned data may provide a geographical criteria basis for decision-making in integrated fruit fly management for guava crops.

## Methods

### Geographical records

Geographic records of *A. striata* were compiled from specimens deposited at following entomological collections: Colección Entomológica de la Universidad de Antioquia, Medellín, Antioquia, Colombia [CEUA], Colección Entomológica de la Universidad Nacional de Colombia, Sede Palmira, Valle del Cauca, Colombia [CEUNP], Colección Taxonómica Nacional Luis María Murillo, ICA Tibaitatá, Mosquera, Cundinamarca, Colombia [CTNI], Colección Entomológica Forestal Universidad Distrital Francisco José de Caldas, Bogotá, Cundinamarca, Colombia [EF-UDFJC], Colección de Insectos del Instituto de Investigación de Recursos Biológicos Alexander von Humboldt, Villa de Leyva, Boyacá, Colombia [IAVH], Colección de Insectos del ICA Palmira, Valle del Cauca, Colombia [ICA-P] Colección de Zoología, Instituto de Ciencias Naturales, Universidad Nacional de Colombia, Sede Bogotá, Cundinamarca, Colombia [ICN], Museo Entomológico “Francisco Luis Gallego”, and Universidad Nacional de Colombia, Sede Medellín, Antioquia, Colombia [MEFLG]; secondary sources, including articles and databases, are also listed (Supp. Table S1). For an adequate geographical interpretation, we provide a plate with the maps of the administrative boundaries (“Departments", as they are locally known in Colombia), natural regions, digital elevation models, and the location of some geographical features referred to troughout the text (Figure 1).

**Figure 1.**
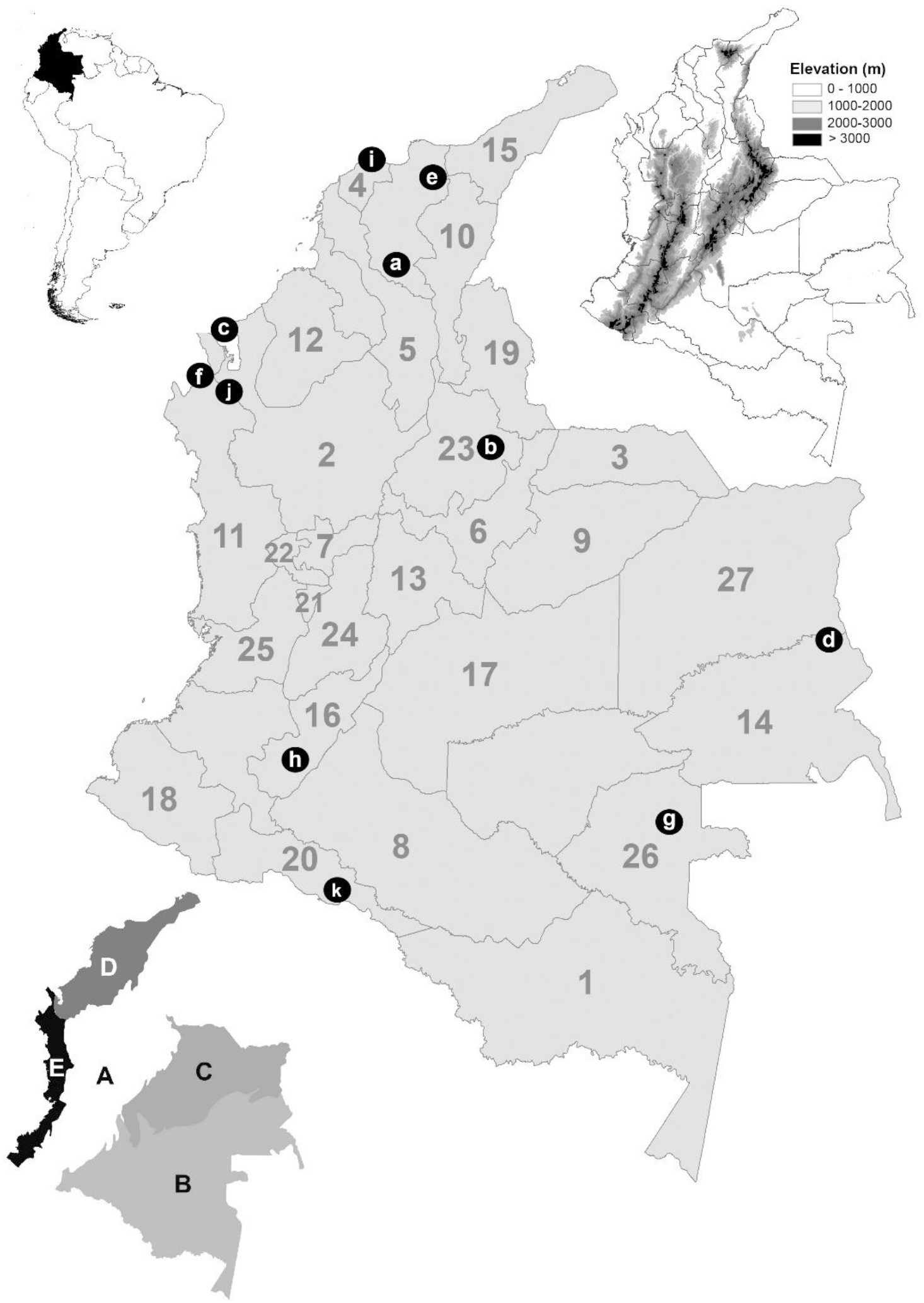
Location of Colombia in South America (upper left). Digital elevation map (upper right) (m) meters. Map of administrative boundaries (center) (number of each department in gray) **1**. Amazonas; **2**. Antioquia; **3**. Arauca; **4**. Atlántico; **5**. Bolívar; **6**. Boyacá; **7**. Caldas; **8**. Caqueta; **9**. Casanare; **10**. Cesar; **11**. Chocó; **12**. Córdoba; **13**. Cundinamarca; **14**. Guainia; **15**. Guajira; **16**. Huila; **17**. Meta; **18**. Nariño; **19**. Norte de Santander; **20**. Putumayo; **21**. Quindío; **22**. Risaralda; **23**. Santander; **24**. Tolima; **25**. Valle del Cauca; **26**. Vaupes; **27**. Vichada. Referenced localities are indicated with black dots and white letters. **a**. Depresión Momposina; **b**. Cañon del Chicamocha; **c**. Golfo de Uraba; **d**. Inirida interfluvial region; **e**. Sierra Nevada de Santa Marta; **f**. Serrania del Darien; **g**. Vaupes River; **h**. Valle de Laboyos; **i**. Salamanca National Park; **j**. Los Katios National Park; **k**. La Paya National Park. Five natural regions (bottom left) **A**. Andes; **B**. Amazonas; **C**. Orinoquia; **D**. Caribbe; **E**. Pacific.

### Climatic information

Bioclimatic variables were gathered from the WorldClim 1.4 climate data archive (Hijmans et al. 2005) (Table 1) in the form of 19 bioclimatic data layers, summarizing potentially relevant climate dimensions at a 30 arc-second (∼1 km) spatial resolution. The date were derived from monthly precipitation and temperature values, appropriate to the biological requirements of *A. striata* in terms of temperature, precipitation, and seasonal trends, and extreme or limiting environmental factors (Hijmans et al. 2005). Bioclimatic data layers incorporating global climate change were calculated using a general circulation model (GCM) for different scenarios. The MIROC5 Global Climate model was selected to include variation and uncertainty among climate change mathematical simulations (Yañez-Arenas et al. 2016). We considered 2050 and 2070 as future time slices, under two emission scenarios: Representative Concentration Pathways (RCPs) 2.6 and 8.5. They are consistent with a wide range of possible changes in future anthropogenic (i.e., human-caused) greenhouse gas (GHG) emissions, and aim to represent their atmospheric concentrations in different scenarios (Ward et al., 2012). RCP 2.6 assumes that global annual GHG emissions (measured in CO2 equivalents) peak will between 2010 and 2020, with emissions declining substantially after that (Meinshausen et al., 2011). Under RCP 8.5, emissions continue to rise throughout the 21st century (Meinshausen et al. 2011).

**Table 1.** Bioclimatic variables used in modeling the potential distribution for *Anastrepha striata* in Colombia.

| Code | Environmental variable | Percent contribution | N° of correlated variables |
| --- | --- | --- | --- |
| Bio1 | Annual Mean Temperature | 4.9 | 3 |
| Bio3 | Isothermality (BIO2/BIO7) (* 100) | 23.3 | 2 |
| Bio4 | Temperature Seasonality (standard deviation *100) | 7.5 | 4 |
| Bio5 | Max Temperature of Warmest Month | 6.5 | 3 |
| Bio6 | Min Temperature of Coldest Month | 13.3 | 4 |
| Bio11 | Mean Temperature of Coldest Quarter | 5.1 | 1 |
| Bio12 | Annual Precipitation | 21 | 2 |
| Bio14 | Precipitation of Driest Month | 5.8 | 3 |
| Bio15 | Precipitation Seasonality (Coefficient of Variation) | 8.3 | 1 |

An essential element in the development of ecological niche models is the hypotheses of areas (M) that have been accessible to the species (Barve et al. 2011). Based on the presence records and the terrestrial ecoregions of the world proposed by the World Wildlife Foundation (Olson et al. 2001), we estimated the area M to calibrate the model. Charted colombian administrative boundaries (Fig. 1) were used as the area for the final projection model. According to the variable contributions calculated by the jackknife analysis and the Pearson correlation coefficients, we determined which variables would be ratained for further evaluations. If two variables had a correlation of > [0.8], the highly contributing variable was preferred over the other (Raghavan et al. 2019). In the current and future models, we used a total of nine bioclimatic variables (Table 1).

### Model design

The potential distribution model was generated with a maximum entropy algorithm incorporated in MaxEnt v.3.3.3k (Phillips et al. 2006). Partial receiver operating characteristic (pROC) statistics were applied for only the current model to the 50% subset of occurrences left out before model calibration for testing. We chose pROC as a significance test in light of critiques of the appropriateness of traditional ROC approaches (Peterson et al. 2008). This metric was used to test the statistical significance of ecological niche model predictions. A value of 1.0 was equivalent to the performance of a random classifier. These results were based on 100 bootstrap replicates, and statistical significance was assessed by bootstrapping and comparison with a random classifier ratio of 1.0 according to the significant sensitivity of this algorithm to particular parameter settings. We conducted a detailed model selection exercise, using the ENMeval R package. This provided an automated method to execute MaxEnt models across a user-specified range of regularization multiplier (RM) values and feature combinations (FCs) (Muscarella et al. 2014). We set the RM range from 0.5 to 4.0, with increments of 0.5, and employed three FCs, i.e., linear (L); linear and quadratic (LQ); linear, quadratic and product (LQP); linear, quadratic, product and threshold (LQPT); linear, quadratic, product threshold and hinge (LQPTH), resulting in 45 possible combinations of features and regularization multipliers (Muscarella et al. 2014). The fine-tuned MaxEnt models were made by seeking the lowest delta value of Akaike’s information criterion, which was corrected for small sample sizes (AICc) among the candidate models, reflecting both model goodness-of-fit and complexity to provide the most conservative results (Basanta et al. 2019). We selected a model with the lowest delta AICc score, which had a parametrization of regularization multiplier of 2.0 and an LQHP feature combination; it exhibited good predictive performance.

A total of ten model replications were implemented through bootstrapping tools. The medians were used through repetitions as a final niche estimation (Altamiranda-Saavedra et al. 2017). All models were converted to binary using a threshold of training omission rate with an error rate of E = 5%. The threshold selection methods were based on lower threshold values, i.e., with a broader distribution of suitable habitat and close to zero errors of omission. To predict variations in the spatial distribution, the expansion and contraction in the dispersion area were estimated through pairwise ranking between the two binary distribution models (current and future distribution models) through the SDMtoolsbox tool in ArcGIS 10.3. Finally, we calculated the range of median values across all models for RCP 2.6 and 8.5, and we considered the estimated variance among models as a measure of uncertainty using ArcGis 10.3 (Peterson et al. 2018). A variance partitioning approach was used to compare the estimates of environmental suitability in the SDM prediction maps on a pixel-by-pixel basis across different maps and to characterize the proportion of variance in the estimates of suitability attributable to individual factors (Diniz-Filho et al. 2009). As supplementary material, all models are available to download in the .*KMZ* format (Supplementary Material 2).

### Current risk of establishment of *A. striata* in municipalities

A preliminary list of 75 guava producer municipalities was generated by consulting annual reports from the ICA (Instituto Colombiano Agropecuario) Phytosanitary Surveillance and Epidemiology Technical Division (ICA 2020). The area at risk for *A. striata* establishment was measured as the percentage of suitable space in the current potential distribution model by each municipality using ArcGIS 10.3. Consequently, the seventy-five districts were classified in the following way: units with coverage below 33% were considered low vulnerability; those with coverage between 33% and 66% had intermediate vulnerability; and those with coverage above 66% had high vulnerability.

## Results

We collected a total of 211 geographical records of *A. striata* at elevations ranging from 6 to 3,044 m (Figure 2a); and most were located in the Andean region (Figure. 1A). The percentage contribution of the bionomical variables is shown in Table 1. The final model for current environmental conditions showed an extensive suitable area for *A. striata* establishment, mainly in the Andean region (Figure 1A). There was less establishment risk in the Caribbean (Figure 1D) and Pacific region (Figure 1E), and almost none existent risk in the Orinoquia (Figure 1C) and Amazonian regions (Figure 1D) (Figure 2a). Despite the notable absence of suitable areas in the Amazonia region, the current model (Figure 2a) included the interfluvial areas of the Inírida, Guainía, and Vaupés rivers (Figure 1. Localities d,g); the surroundings jurisdiction of Mitú in Vaupes (Figure 1.26); southwestern Putumayo (Figure 1.20); and the western area of La Paya National Natural Park (Figure 1. locality k) as suitable for *A. striata* establishment. Glaring errors of omission were evidenced in locations such as Leticia in southern Amazonas (Figure 1.1); southwestern Putumayo (Figure 1.20); Orinoquian localities, such as northeastern Vichada (Figure 1.27), and northern Arauca (Figure 1.3); and finally in the Caribbean in central Guajira (Figure 1.15). The currently unsuitable areas for the establishment of *A. striata* (Figure 2a) were as follows: in the Caribbean region, a large part of the xerophytic formations in northern Guajira (Figure 1. 15), areas of Salamanca National Park (Figure 1. Locality i), the Depresión Momposina (Figure 1. Locality a), Sierra Nevada de Santa Marta (Figure 1. Locality e) and swamp complexes in eastern Cordoba (Figure 1.12). Areas above the 2,050 m elevation along the Andean region were also unsuitable, including all high altitude areas of the Andean paramos complex (Figure 1A); the foothills of western Norte de Santander (Figure 1.19), eastern Boyacá (Figure 1.6) and Cundinamarca (Figure 1.13); and the surrounding areas of El Cañon de Chicamocha (Figure 1. locality b) and Valle de Laboyos in Huila (Figure 1. Locality h). Extensive areas of tropical rainforest (TRF) in the Pacific and the Amazon (Figure 1B, E) were deemed unsuitable, as were the savannas in the Orinoquia region (Figure 1C).

**Figure 2.**
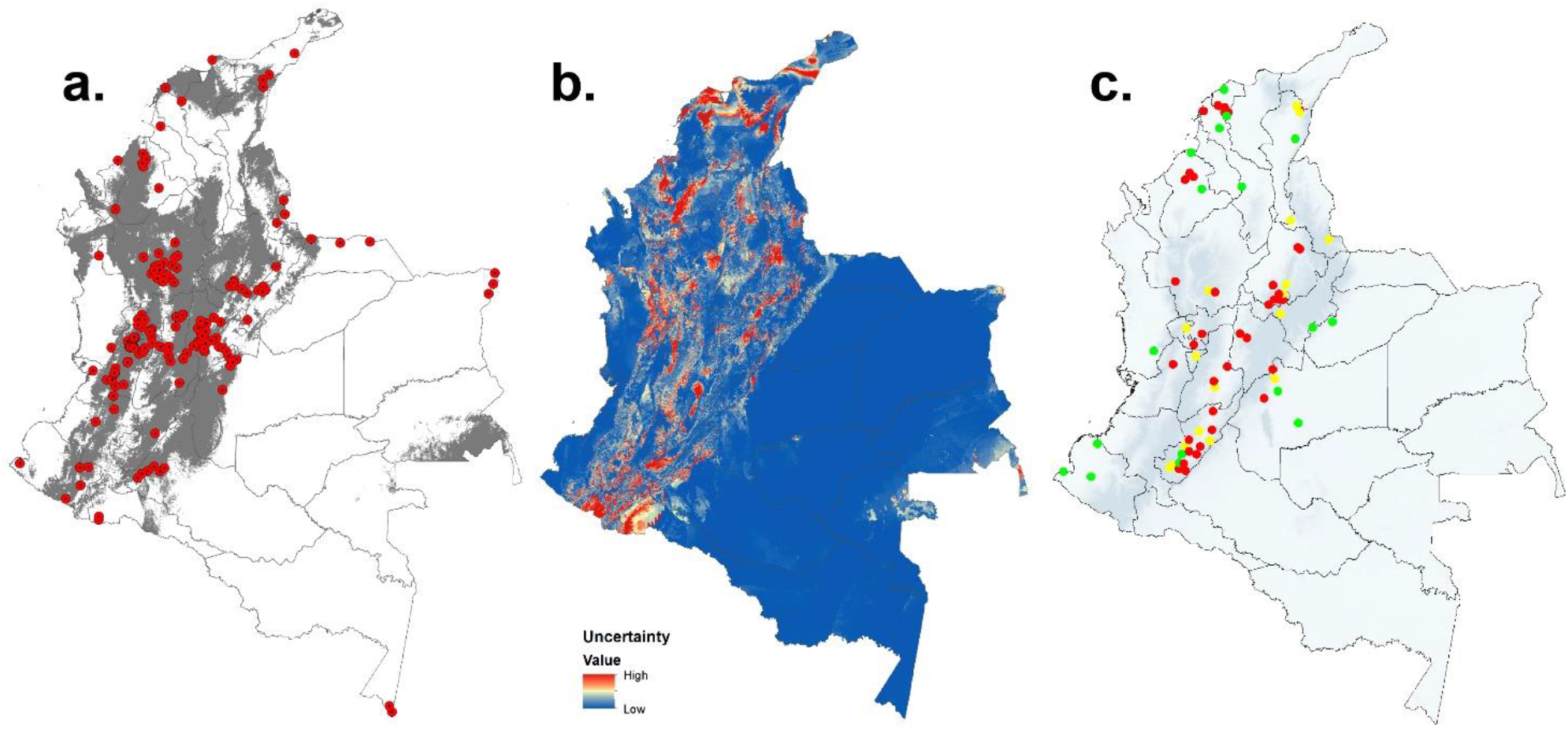
**a)**. Potential distribution of *Anastrepha striata* in Colombia for the current environmental conditions (suitable areas in gray); red dots are the localities of the compiled geographical records; **b)**. Uncertainty of models in the range of median values of general circulation models for *Anastrepha striata*. The color scale represents the degree of variance (blue:low; red:high); **c)**. Locations of guava producer municipalities and their vulnerability category for *Anastrepha striata* establishment under the current climatic scenario. Red) high; yellow: intermediate and green: low.

**Figure 3.**
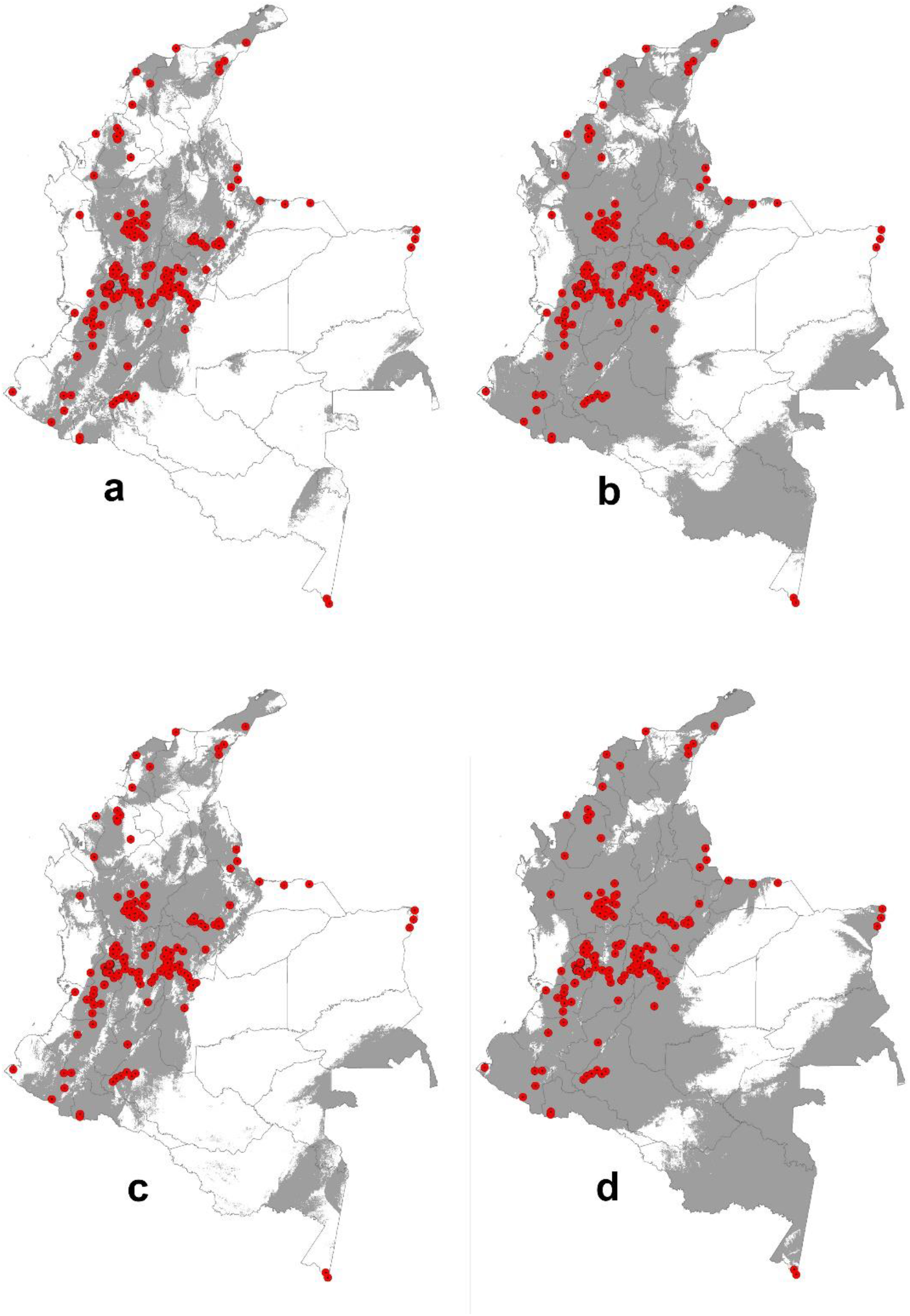
Potential distribution of *Anastrepha striata* in Colombia transferred to climate change scenarios. **a)**. 2050 under Representative Concentration Pathways (RCP) 2.6, **b)**. 2050 under Representative Concentration Pathway (RCP) 8.5, **c)**. 2070 and Representative Concentration Pathways (RCP)2.6, **d)**. 2070 Representative Concentration Pathway (RCP) 8.5.

### Potential distribution under climate change scenarios

Significant levels of uncertainty in climate change models were found, specially in the Andean region (Figure 3). An increase in the area suitable for *A. striata* establishment was observed in all climate change scenarios (Figure 4a,b,c,d). This result was more pronounced in the most pessimistic greenhouse gas emission scenario (RCP 8.5), for both temporal projections in 2050 and 2070 (Fig. 4B and 4D). According to the analysis of predicted changes in suitable habitat and the associated potential distributions, the greatest extent of the possible area increase for *A. striata* was predicted under the RCP 8.5 scenario by 2050 (Figure. 5b) with an increased area of 520,071 km^2^ (Table 2). Remarkably, this expansion was predicted to occur mainly in the Amazonian natural region (Figure 1B), including the departments of Caquetá (Figure 1.8), Amazonas (Figure 1.1), Vaupés (Figure 1.26), Guainía (Figure 1.14), and Putumayo (Figure 1.20). On the other hand, RCP 2.6 predicted reductions in the area (i.e., contraction area) by 2050, with a potential decrease of more than 52, 808 km^2^ (Table 2). This effect was specially distinct in the northern area of the Pacific region, specifically in the Chocó department (Figure 1.11 and 4a).

**Table 2.** Changes in the potential distribution area (km^2^) of *Anastrepha striata* in Colombia between different climatic scenarios.

| Model | Expansion | Absence in both | Presence in both | Contraction |
| --- | --- | --- | --- | --- |
| Current/MIROC5-2050-RCP26 | 98,611 | 730,745 | 253,462 | 52,808 |
| Current/ MIROC5-2050-RCP85 | 369,239 | 460,118 | 297,437 | 8,834 |
| Current/ MIROC5-2070-RCP26 | 187,860 | 641,497 | 265,375 | 40,895 |
| Current/ MIROC5-2070-RCP85 | 520,071 | 309,285 | 303,298 | 2,972 |

**Figure 4.**
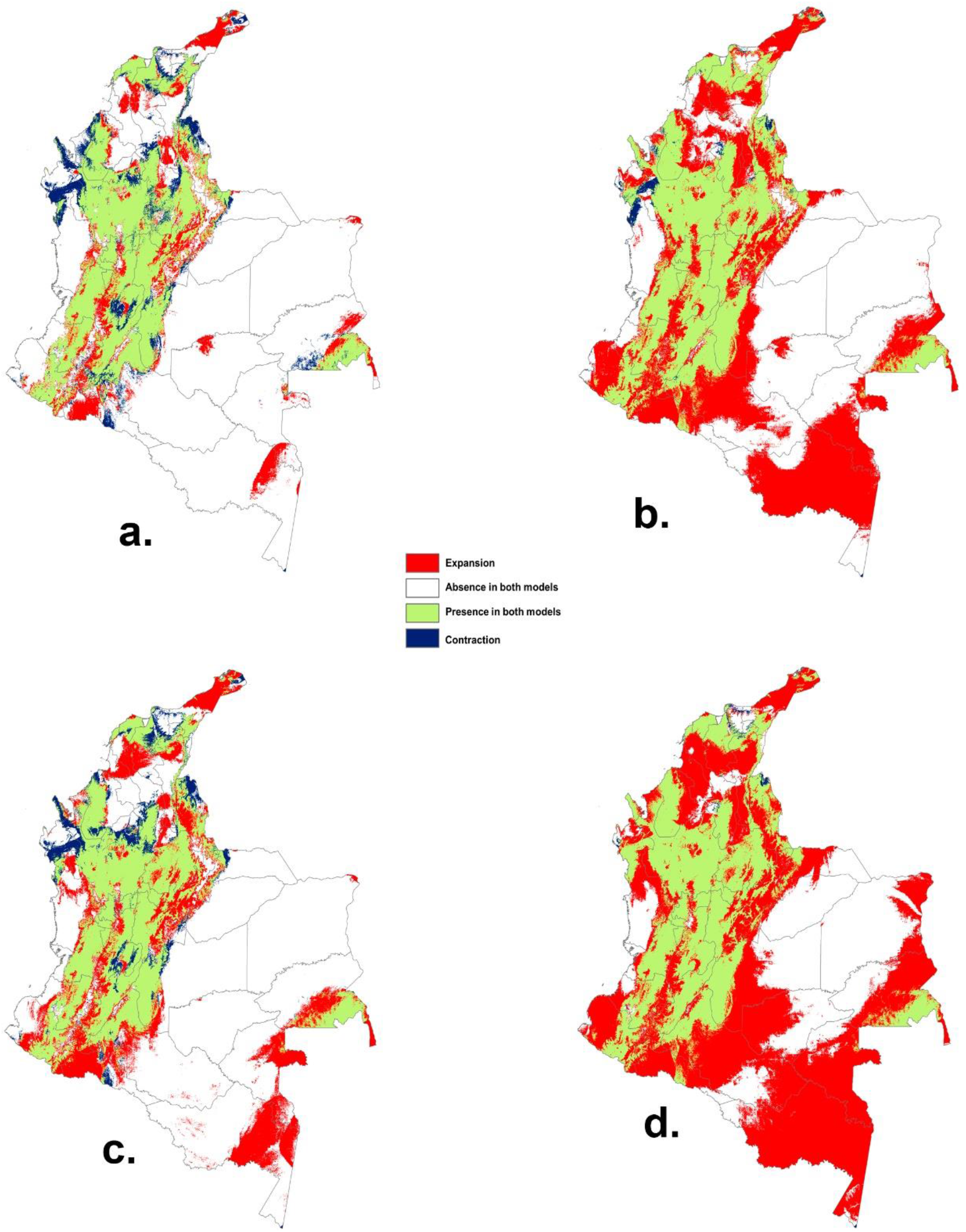
Changes in suitable climatic spaces for *Anastrepha striata* and the potential distributions between current and future conditions in Colombia, **a)**. Current vs. time 2050 under RCP 2.6, **b**.) Current vs. 2050 under RCP 8.5 **c**). Current vs. 2070 under RCP 2.6, **d**.) Current vs. 2070 under RCP 8.5.

### Current risk of establishment of A. striata in municipalities

Forty-eight guava-producing municipalities in Colombia are located in the Andean region (Fig. 1A), 18 in the Caribbean region (Fig. 1D), 8 in the Pacific region (Fig. 1E), and 6 in the Orinoquia region (Fig. 1C) (Figure 2c). Of these municipalities, 56 % were categorized as highly vulnerable to the establishment of *A. striata*, 21 % had intermediate vulnerability, and 23% had low vulnerability (Figure 2c and Table 3).

**Table 3.**
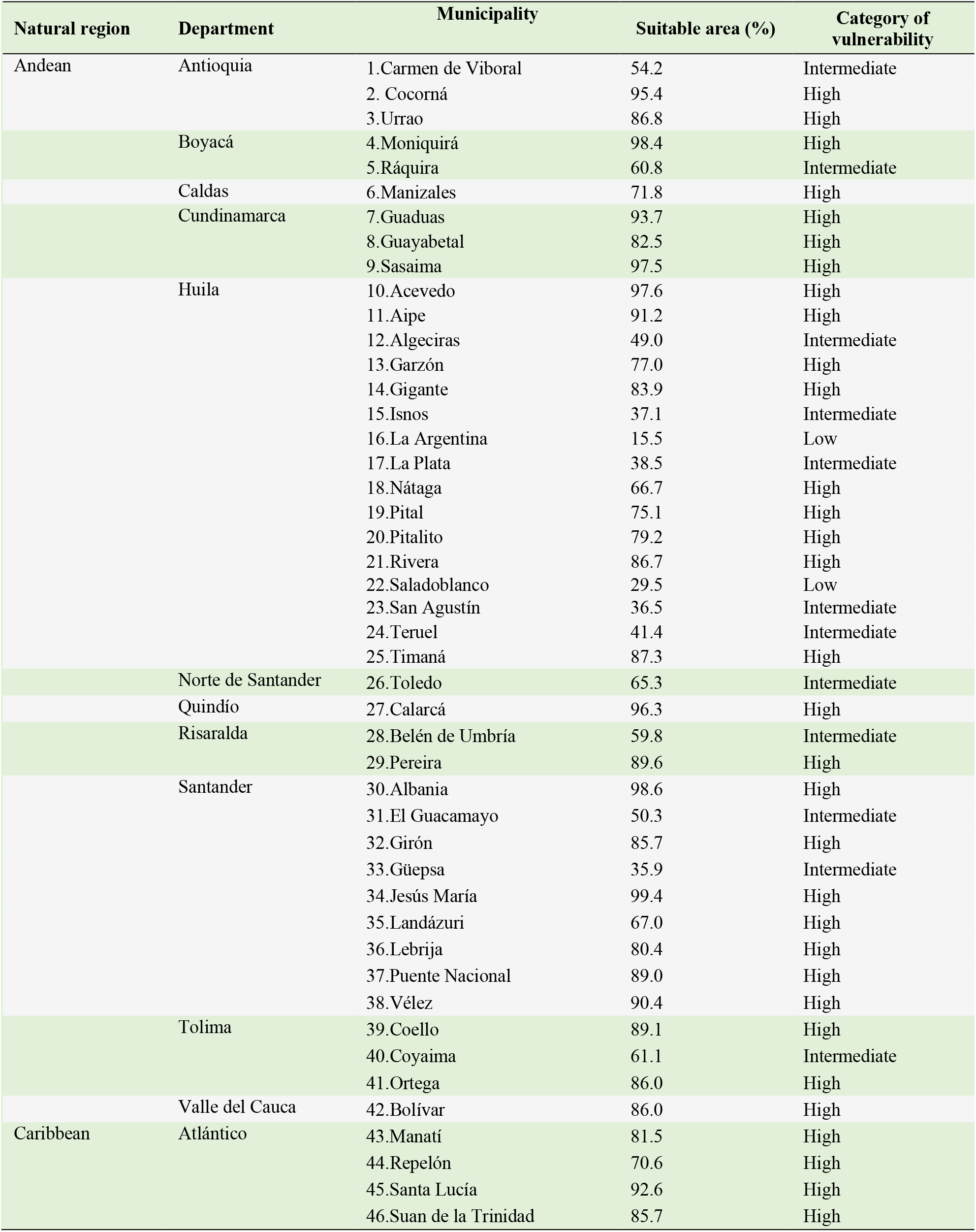
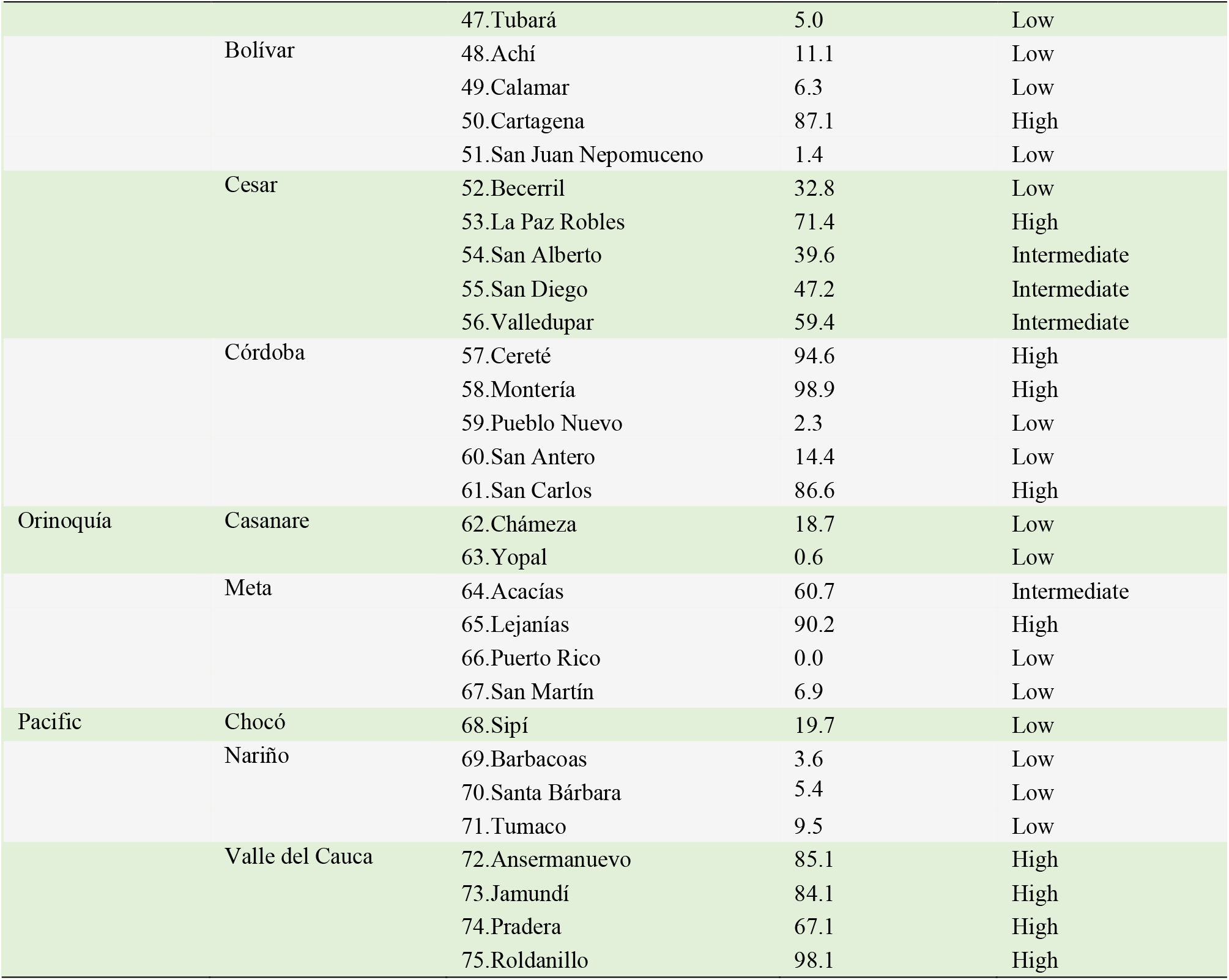
List of guava-producing municipalities and their respective risk category for the establishment of *Anastrepha striata* according to the area deemed suitable by the ecological niche model under current climate conditions in Colombia.

## Discussion

This study is the first regional (northwest South America) approach tho use ecological niche modeling to assess the potential distribution of a fruit fly species of economic importance under global climate change scenarios. Aditionally, this study is the first to consider the risk of pest establishment according to administrative boundaries to configure regional policies and decision making. The results showed that in Colombia, i) under the current environmental model, the suitable areas for the establishment of *A. striata* were located mainly in the Andean region, with some risk area in the Caribbean region and to a lesser extent in the Pacific, Orinoquia and Amazonian regions. High spatial alignment with geographical records was recently reported by Rodriguez Clavijo et al., (2018). ii) The climate change models showed an increase in suitable areas for the establishment of the *A. striata* in response to global warming, and iii) high environmental suitability for the establishment of populations was evidenced in the guava-producing municipalities in Colombia.

The retrieved geographical records of *A. striata* were mainly based on two sources: the first came from five departmental initiatives (control programs) (Castañeda et al. 2010) and the second came from sporadic records of specimens deposited in entomological collection. These data demostrated the lack of comprehensive monitoring of fruit fly management and a rigorous systematic phytosanitary surveillance program. Optimal biological data are a crucial aspect for good results in mitigating the effects of the fly, and avoiding duplication of efforts; different Colombian environmental agencies operate under varying policies. Despite these circumstances, the current distribution model coincided with previous information, which showed that guava fruit flies were common in Colombia, as summarized by Rodriguez Clavijo et al., (2018). The distribution of fruit fly species in Colombia is also related to the presence of its primary host plant (Castañeda et al. 2010). *A. striata* is associated mainly with Myrtaceae, a family encompassing nearly 180 species in Colombia that is distributed across all vegetation types communities and altitudinal gradients (Parra-0 2014).

Additionally, it is commonly found among the most diverse plant families in vegetation surveys, either in the Amazonian region or in the ecotones of Páramo (Parra-0 2014). However, the prevalence of *A. striata* in Colombia may be associated not only with the presence of its host plants but also with favorable environmental conditions or factors that regulate their trophic relationships (Hedström 1991). Our results indicated that *A. striata* currently inhabits a significant portion of the Colombian territory, not only because of the above mentioned factors but also because of its thermal physiological plasticity (Baker et al. 1944) related to the climate along the Andean altitudinal gradient (between 6 to 3,044 m. elevation, ranging from 9.2°C to 23.8°C). Low misture or high or low temperature regimes may pose physiological restrictions for both the fly and its hosts. This fact was evidenced by Stone (1939), who reported high mortality rates in *A. striata* larvae exposed momentarily to temperatures of 40°C. Similarly, Bolzan et al. (2017) reported no embryonic development in regimes above 35°C.

The most influential variables of *A. striata* occurrence were the temperature and the precipitation-related variables (Table 1); these results aligned with those of Porter et al., (1991), who reported that these factors significantly affected the distribution of pest insects. The models also indicated absence of the flies in cold localities at high altitudes, such as paramo ecosystems; the paramo is distributed along the top of the Andean ranges (Figure 1A) and the upper Sierra Nevada de Santa Marta (Figure 1 locality e). Additionally, very humid regions with high precipitation levels (tropical rainforests) such as the Serrania de Darien (Figure 1 Locality f) and most forested areas of the Pacific (Figure 1E) and Amazonia region (Figure 1B) were identified as having low suitability for the flies. It has been showed that less disturbed ecosystems, such as tropical rainforests, offer fewer host plant species, lower infestation rates, and more ecological complexity, which probably reduces fruit fly pest performance (Hernández-Ortiz and Pérez-Alonso 1993). Eventually, the land use in these areas, which are considered to have high forestry potential, may result the fruit fly introduction (Mena et al. 2015). The area suitable for *A. striata* in the Pacific region was mainly located in northern Chocó (Figure 2a. and Figure 1.11), in extensive areas bordering the Golfo de Uraba (Figure 1. Locality c) and in a small area in the Nariño department (Figure 1.18). The low suitability of habitat in this natural region might be associated with the forest conservation levels and the vocation for the agroforestry industry of the region (De la Hoz 2007). However, the map indicates suitability in the interfluvial area of the Inírida (Figure 1 locality d), Guainía and Vaupés rivers, and the surroundings of the jurisdiction of Mitú (Figure 1 locality g) in the Orinoquia and Amazonian regions. The establishment of *A. striata* must be interpreted cautiously since the precarious conditions of the soils, where it is common to find rocky outcrops and floristic associations of monocotyledons (Hernández-Camacho and Sanchez-Páez 1992), provide unfavorable conditions for *A. striata*’s host plants. However, bionomic and scenopoetic variables not assessed there may positively affect the occurrence of source populations (Peterson et al. 2011).

The models under climate change scenarios with an increase in temperature, expanded the geographic area suitably, as evidenced here for *A. striata* in Colombia (Figure 4). This behavioral response may be evidenced in species limited by low temperatures, where the increase in warmth in the occurrence area may shift the geographical range towards cold regions (Fu et al. 2014). Although the presence probability decreased in the Pacific region, it could be the result of climatic effects due to proximity to the coastal zone, where the general climatic conditions are remarkably unstable (Martínez-Ardila et al. 2005). Changes in climatic variables, such as precipitation regimes, can cause contraction in the spatial distribution (Martínez-Freiría et al. 2016). Our results were in agreement with those of Fu et al., (2014), who demostrated that climate change expanded the potential distribution of the fruit fly *Anastrepha obliqua* (Macquart, 1835) towards the poles but decreased the distribution in northwestern Australia and northern sub-Saharan Africa due to climate stress caused by marine climate effects.

It should be noted that the potential distributions of species depend not only on weather conditions but also on dispersal capacity, host availability, and the effects of ecological relationships (Peterson et al. 2011). The estimation of these aspects is especially critical for species of economic importance (Lira-Noriega et al. 2013). This information is difficult to model with climate change scenarios, and even current biotic interaction data are challenging to include (Peterson et al. 2011). This study faced a poor understanding of the basics bionomic parameters of *A. striata* (Cruz-López et al. 2015) due to insufficient local data to infer ecological and distributional patterns in *A. striata* populations (Canal 2010; Castañeda et al. 2010). The current and future ecological interactions of *A. striata* in Colombia are still enigmatic and lead to additional challenges for integrated management.

Nevertheless, we offer an additional tool never before considered in Colombian fruit agriculture. The most significant proportion of potential areas predicted by climate change scenario models for the expansion of *A. striata* occurred in the Amazonian region (Figure 4b,d). This result could be related to accelerated deforestation rates, which are caused primarily by the presence of illicit crops and expansion of the agricultural frontier (Vieira 2019); these activities could promote *A. striata* establishment (Aluja et al. 2003).

Due to the economic importance of *A. striata*, knowledge on the autoecology and variables determining its geographic distribution is essential at the local scale (Castañeda et al. 2010); this information provides crucial feedback for implementing effective integrated pest management programs (IPMs (Martínez-Ardila et al. 2005). Our results indicated that under the current environmental and climatic conditions, the *A. striata* distribution is intimately associated with guava crops. The predominance of *A. striata* and guava crop interactions, the high vulnerability of the guava producer municipalities to the potential occurrence of *A. striata*, and its presence in a large area principally on the Andes (Figure 2c, Table 3) make it difficult to effectively establish integrated pest management strategies based on a single local initiative. We encourage the use of the offered data here concerning each municipality to configure national policies based on area-wide management (AWM).

Furthermore, the estimated distributions for *A. striata* according to climate change scenarios for 2050 and 2070 will not modify this outlook and are trending towards expansion. Importantly, the data presented here have established a clear and present risk to the spread of this fly of economic importance, emphasizing that these risks will only worsen in the face of climate change. Action needs to be taken to ensure optimal guava productivity. The Selection and cultivation of cultivars adapted to environments unsuitable for *A. striata*, as well as the selection of fly resistant varieties, present promising opportunities. Alternative approaches employing chemical ecology and trophic relationship studies could represent useful improvements for guava fruit fly management in Colombia. Finally, it would be desirable to establish transnational policies to enhance monitoring of fruit fly pests in areas where eradication techniques, such as low prevalence areas and sterile individulas use, are unlikely.

## Acknowledgments

We thank Javier Martínez for allowing us to access to the *A. striata* distribution records, Juan Manuel Perilla for information compilation, and the curators of the referenced entomological collections. This work was supported by Tecnológico de Antioquia (CODEI), and COLCIENCIAS, who financed Juan Manuel Perilla as a young researcher. The authors disclose that they do not have any financial and personal relationship with other people or organizations that could inappropriately influence (bias) this work. The authors also state that they have no conflicts of interest.

